# The BioImage Archive - building a home for life-sciences microscopy data

**DOI:** 10.1101/2021.12.17.473169

**Authors:** Matthew Hartley, Gerard J. Kleywegt, Ardan Patwardhan, Ugis Sarkans, Jason R. Swedlow, Alvis Brazma

## Abstract

Despite the huge impact of data resources in genomics and structural biology, until now there has been no central archive for biological data for all imaging modalities. The BioImage Archive is a new data resource at the European Bioinformatics Institute (EMBL-EBI) designed to fill this gap. In its initial development BioImage Archive accepts bioimaging data associated with publications, in any format, from any imaging modality from the molecular to the organism scale, excluding medical imaging. The BioImage Archive will ensure reproducibility of published studies that derive results from image data and reduce duplication of effort. Most importantly, the BioImage Archive will help scientists to generate new insights through reuse of existing data to answer new biological questions, and provision of training, testing and benchmarking data for development of tools for image analysis. The archive is available at https://www.ebi.ac.uk/bioimage-archive/.

**Highlights:**

- The BioImage Archive is a new archival data resource at the European Bioinformatics Institute (EMBL-EBI).
- The BioImage Archive aims to accept all biological imaging data associated with peer-reviewed publications using approaches that probe biological structure, mechanism and dynamics, as well as other important datasets that can serve as reference examples for particular biological or technical domains.
- The BioImage Archive aims to encourage the use of valuable imaging data, to improve reproducibility of published results that rely on image data, and to facilitate extraction of novel biological insights from existing data and development of new image analysis methods.
- The BioImage Archive forms the foundation for an ecosystem of related databases, supporting those resources with storage infrastructure and indexing across databases.
- Across this ecosystem, the BioImage Archive already stores and provides access to over 1.5 petabytes of image data from many different imaging modalities and biological domains.
- Future development of the BioImage Archive will support the fast-emerging next generation file formats (NGFFs) for bioimaging data, providing access mechanisms tailored toward modern visualisation and data exploration tools, as well as unlocking the power of modern AI-based image-analysis approaches.

## Introduction

Imaging is a key research tool in the life sciences^1,2^. “Biological imaging” encompasses a broad range of modern microscopy and other methods that generate large and complex datasets. It comprises a diverse set of subdomains providing spatial and temporal information on biological systems and their components at different physical scales. In most modalities, biological imaging involves a combination of data acquisition and nontrivial data-analysis workflows that together provide results that can be interpreted by biologists.

Such data creates significant opportunities for reuse, potentially leading to new scientific discoveries as well as maximising the return on investment in data generation. To enable this reuse, access to open image data that follows the FAIR principles^3^ is a prerequisite. Where such access has been provided for sequence and structural data, through key resources such as the ENA^4^ and PDB^5^, the scientific impact has been immense. Biological data resources have become a critical part of the infrastructure for life-sciences research^6^. Bioinformatics databases can be broadly categorised into either deposition databases (archives) or added-value databases^7^. Deposition databases create a persistent scientific record of the data on which published scientific conclusions are based, by providing deposition pipelines for data and associated metadata and making that data searchable and accessible to the community. Added-value databases enrich data through expert curation, data integration and further analysis, often across multiple datasets, each individually archived.

Over the past decade, several resources have emerged that have begun to tackle the challenge of publishing bioimaging datasets. In 2013, EMBL-EBI launched EMPIAR, the Electron Microscopy Public Image Archive, in response to community demand^8^ for public archiving of raw 2D image data to support the validation of 3D cryo-EM structures^9^. In 2016, a collaboration between the OME consortium and EMBL-EBI resulted in the Image Data Resource (IDR), a platform for bioimage data integration and reanalysis^10^. In parallel, the Systems Science of Biological Dynamics Database (SSBD)^11^ began publishing biological imaging datasets that captured temporal changes in various model systems. In addition, several institute- or project-specific resources have emerged that make cell or tissue imaging datasets available (for example, the Allen Cell Explorer^12^, or the NCI Imaging Data Commons^13^). Although these added-value resources met specific demands, there remained a gap in provision for a broader image archive, leading to a community call for the development of a public bioimage archive in 2018^14^.

In 2019, EMBL-EBI launched the BioImage Archive to begin to address this need. The initial launch provided both a central hub to link together IDR and EMPIAR, and direct data deposition through BioStudies, EMBL-EBI’s resource for data integration and data that does not fit into existing specialised archives^15^. The BioImage Archive now operates as a data resource of its own identity, separate from its historic antecedents. It supports rapid direct deposition of novel imaging datasets associated with publications, import of datasets from other biological imaging resources, and integration with other data resources.

## Results

### Purpose and initial focus

The primary goals of the BioImage Archive are to:

1. Provide a single home for biological imaging data and facilitate the discoverability of these data.
2. Maximise the use of imaging data that can be expensive or even impossible to acquire repeatedly (for example if the imaged sample was unique).
3. Ensure the reproducibility of scientific results based on biological imaging.
4. Enable new discoveries to be made, and insights to be gained, from existing data by encouraging their reuse.
5. Accelerate the development of image analysis methods.

To achieve these goals, the archive needs to provide access to a wide range of bioimage data in a way that facilitates discoverability and reuse, as the FAIR principles set out^1^. This requires supporting straightforward direct deposition of new data, indexing, search and retrieval of datasets in reuse and visualisation-friendly formats, and integration with other data resources. To address the immediate demands of the community, while building infrastructure and processes to support long-term growth and wide data reuse, the initial focus of the archive is on:

a. Providing a rapid and straightforward deposition process, such that submitters preparing for publication can receive an accession identifier for the imaging data upon publication.
b. Building a diverse collection of image data, while steadily improving the quality of associated metadata.
c. Enhancing reusability of those images through discoverability based on rich metadata and easy access to both whole datasets and individual images.
d. Supporting added-value databases through resource indexing, provision of storage infrastructure, data import/export and linking between resources.

### Data deposition and scope

The BioImage Archive accepts open biological imaging data associated with publications from any imaging modality, at scales from Ångströms to centimetres. The archive also accepts “reference” image datasets, where data clearly provide value beyond a single experiment or study. It cannot, however, accept patient-identifiable medical data, such as that derived from clinical imaging.

### Submission process

The BioImage Archive provides a lightweight submission process, with four stages^16^:

1. Preparation for submission - organising data and registering an account if needed.
2. Upload of data and preparation of file-level metadata.
3. Completion of dataset-level metadata.
4. Finalisation of submission.

Submitters organise their data locally before submission, by creation of a suitable file and directory structure. The BioImage Archive allows considerable flexibility in the structure of submitted datasets in order to allow the archive to support the wide range of imaging modalities and experimental setups, as well as to allow submission of intermediate and downstream analysed data for which no general formal structure yet exists.

Submitters then upload their image and supporting data files. For smaller datasets, this can be done through the submission tool, which provides a graphical interface for file uploads. For larger datasets, FTP and Aspera uploads are supported. These large datasets may take several days to upload, so planning depositions ahead of time is recommended, particularly if publication is dependent on data release. Submitters receive a unique tracking identifier for their submission so that they can return to their deposition session.

They then complete a web form to supply metadata about the images and associated data. This metadata includes information about the submitting team, the study-associated publication that the image data supports, experimental and imaging protocols and summary information about the images themselves. After submission, a permanent accession identifier is assigned. Submission can take place in as little as one day, though the limiting factor is usually data-transfer time.

Some studies generate data that are submitted to multiple resources, such as correlative imaging involving 3D electron (EM) and 3D light microscopy (LM). In this example, EMPIAR accepts the 3D EM data, while the BioImage Archive accepts both the LM data and information on how to link the components together, such as physical transformations into a common coordinate space. A more detailed description of how the BioImage Archive and EMPIAR together support deposition of correlative datasets that involve both resources is described in detail in a separate protocol paper^16^.

### Access, downloads and use

The BioImage Archive provides a web-based interface to browse, search, view metadata and retrieve datasets. This portal presents the dataset-level metadata associated with an accession together with the images and supporting files that accompany that accession.

A flexible search system allows both simple keyword search and construction of complex queries over a range of metadata. When viewing a dataset, information associated with each individual file can be viewed in tabular form. Views can be filtered by this image-file level metadata, allowing relevant subsets of the dataset to be viewed. For example, in a high-content screen, those images associated with a specific compound in the screen can be selected and displayed.

Whole datasets, as well as individual images, can be downloaded either directly through the web interface or via the FTP or Aspera protocols.

### Current usage and growth

At time of writing, the BioImage Archive website is accessed by approximately 2000 different users (measured by unique IP addresses) per month, with visitors coming from a wide range of geographical locations. The archive now indexes over 1200 individual datasets across its component resources, which together represent more than 1.5 petabytes of data. Collectively these datasets represent the output of over 900 unique publications. On average, approximately 120 datasets are accessed or downloaded per month at time of writing. Although newly established, a recent global user survey identified the BioImage Archive as one of the EBI’s 20 most used data resources (https://www.ebi.ac.uk/about/our-impact/impact-report-2021).

The archive’s collections are divided into three categories:

1. Datasets deposited directly into the BioImage Archive.
2. Datasets indexed from resources which hold the actual data (EMPIAR).
3. Data imported from other resources, including databases that are now discontinued, for example the Journal of Cell Biology (JCB) DataViewer^18^.

The number of datasets directly deposited to the BioImage Archive is growing rapidly (**Figure 2**).

**Figure 1.**
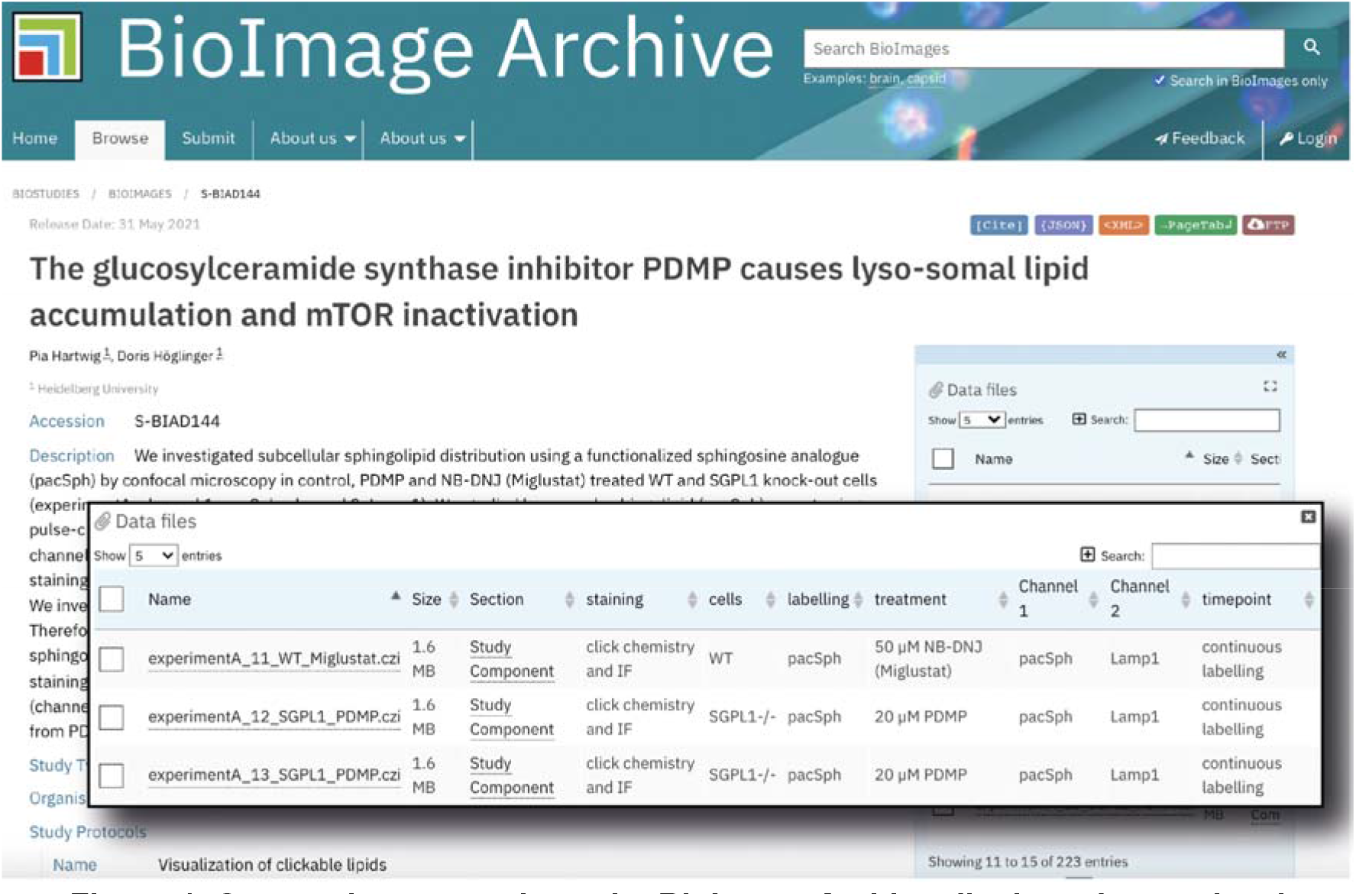
Composite screenshot - the BioImage Archive displays dataset-level metadata (main panel), as well as image-specific metadata (inset). Images can be searched, filtered and downloaded based on this image-specific metadata. Example from S-BIAD144^17^, https://www.ebi.ac.uk/biostudies/BioImages/studies/S-BIAD144.

**Figure 2.**
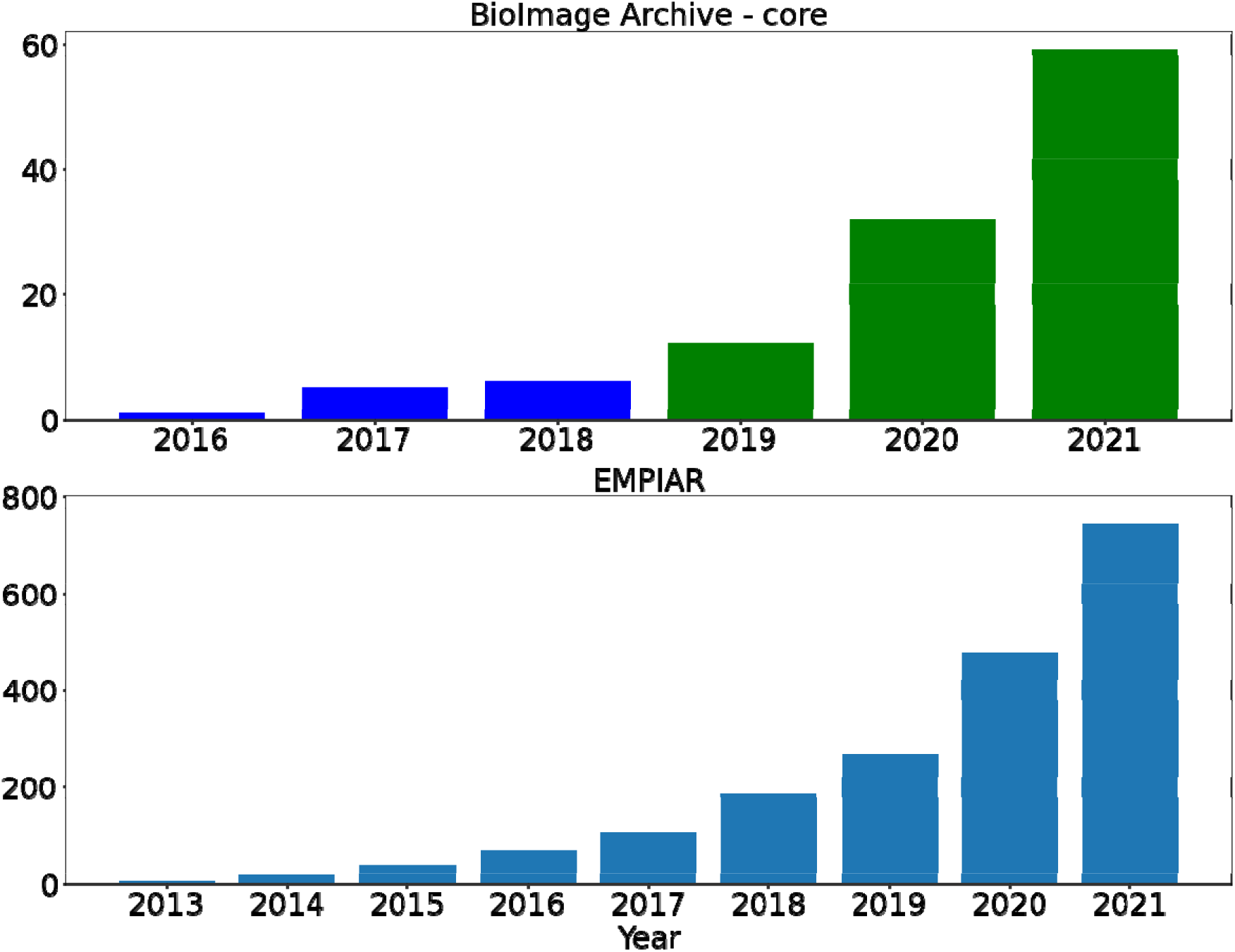
Cumulative numbers of image datasets indexed by the BioImage Archive, up to the end of 2021. In the top panel, the launch of the BioImage Archive is indicated by the colour change of the bars. All JCB datasets were imported in 2018, and their number is constant (424 datasets) and not included in the graphs. The sizes of the datasets vary considerably, from tens of megabytes to dozens of terabytes for a single dataset. The number of individual files comprising a dataset ranges from one to 1.3 million.

As part of its role in underpinning an ecosystem of bioimaging resources, the BioImage Archive indexes data deposited directly to EMPIAR, allowing for inclusive search of those datasets through a single web portal. Data can then be directly viewed and accessed on the EMPIAR pages.

The majority of accessions are from light and electron microscopy. However, as the archive’s broad scope would indicate, many less common technologies including Atomic Force Microscopy (AFM), Micro-CT, and Ultrasound are also represented. By data volume the majority of image files in the core collection of the BioImage Archive are in TIFF format (approximately 75%), but formats reflect the diversity of imaging technologies with over 30 different file types in use.

### BioImage Archive ecosystem and the BioStudies database

Beyond its role in the provision of a service for deposition, indexing and retrieval of biological imaging datasets, the BioImage Archive also supports added-value imaging data resources. The two resources that work most closely with the BioImage Archive are EMPIAR and IDR. We expect that in future other imaging data resources can use this service.

Initially founded to provide a home for raw images underpinning 3D cryo-EM maps and tomograms, EMPIAR has expanded to cover volume EM and 3D X-ray imaging data. In addition to the indexing of EMPIAR datasets described above, the BioImage Archive provides capacious object storage to EMPIAR. As described above, the two resources also work together to support the deposition of correlative imaging data^16^ where different modalities are used to image the same biological specimen.

As mentioned above, IDR is a platform for bioimage data integration and reanalysis. It runs on EMBL-EBI’s Embassy science cloud system (www.embassycloud.org)^19^. The BioImage Archive provides IDR with guarantees of long-term sustainability for data in its collections through import of data from it. Work is also underway to develop a submission mechanism allowing data to be submitted to the IDR through the BioImage Archive, such that submitters can receive an accession identifier quickly for immediate publication of their results, while suitable reference datasets can benefit from the curation and data enrichment provided by IDR.

**Table 1.**
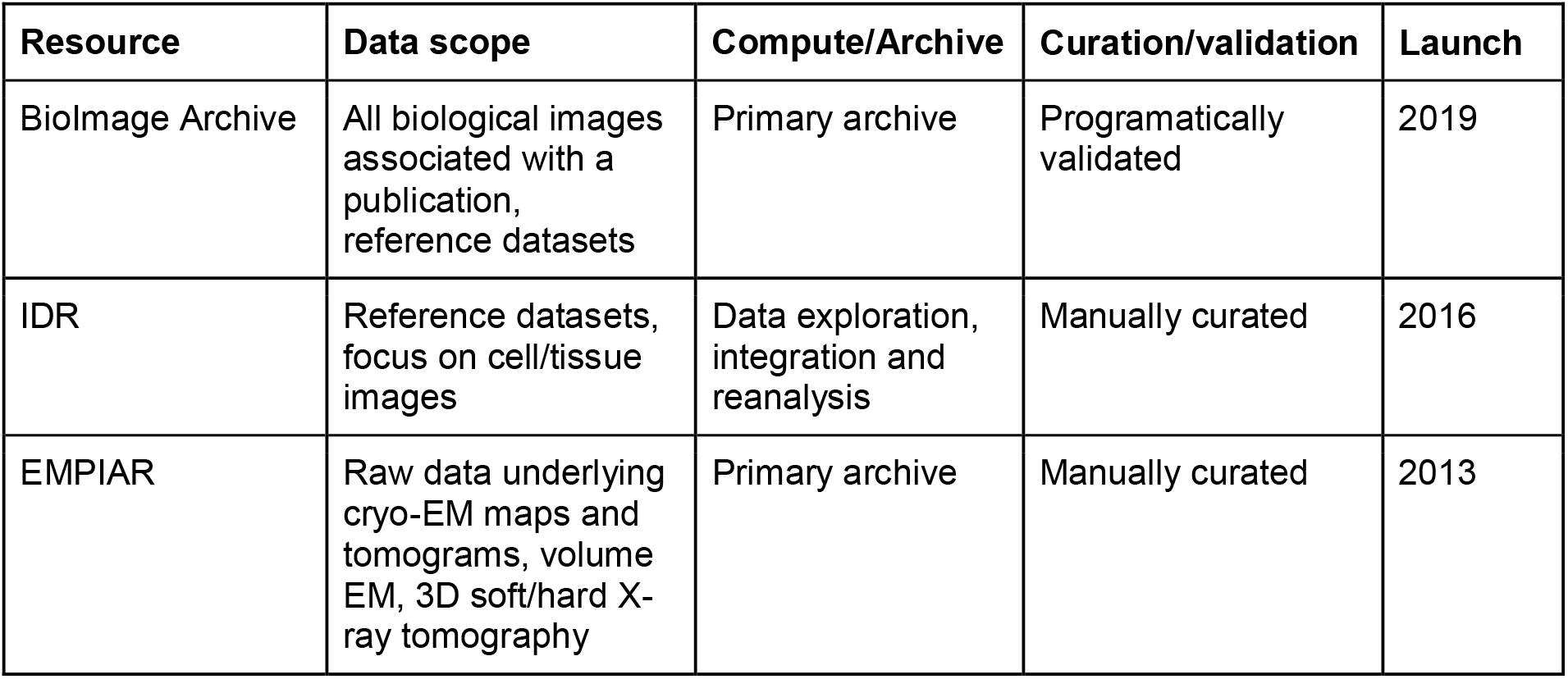
Characteristics of the resources described. IDR provides a platform for computation on its data. IDR/EMPIAR deposition involves manual curation by expert archive staff, while entries deposited into the BioImage Archive undergo an automatic validation process.

| Resource | Data scope | Compute/Archive | Curation/validation | Launch |
| --- | --- | --- | --- | --- |
| BioImage Archive | All biological images associated with a publication, reference datasets | Primary archive | Programmatically validated | 2019 |
| IDR | Reference datasets, focus on cell/tissue images | Data exploration, integration and reanalysis | Manually curated | 2016 |
| EMPIAR | Raw data underlying cryo-EM maps and tomograms, volume EM, 3D soft/hard X-ray tomography | Primary archive | Manually curated | 2013 |

The BioImage Archive builds on this foundation provided by BioStudies, using it as a platform. The BioStudies database at EMBL-EBI enables authors to package all data supporting a publication, through both direct data hosting and integration of links to other data sources^15^. At EMBL-EBI, BioStudies fills a key role by providing a home for data that do not fit into existing dedicated archives. This role enabled BioStudies to develop a pilot service for storage and indexing of image data at EMBL-EBI before the BioImage Archive was established.

## Discussion and Future Plans

As open access archiving of images supporting scientific results increasingly becomes a requirement for publication, and requirements from funding agencies and parent institutes become stricter, the role of the BioImage Archive in providing a service to meet these needs will grow. For both EMPIAR and IDR, major journals such as eLife and Nature Communications began to require deposition to those resources as they became widely used within their communities. Four Cambridge University Press journals, including the microscopy-focussed *Biological Imaging*, recently made the BioImage Archive their recommended deposition repository for imaging data and we expect that Although a young resource, the BioImage Archive is already experiencing rapid growth. Early uptake has been very rapid, with the demand for fast deposition of data to receive an identifier for publication (driven by journal requirements) a notable factor driving submission.

### Current limitations

While the BioImage Archive provides a useful service to data depositors and users, it currently has a number of limitations. Although the archive accepts imaging data from preclinical research, it does not support medical imaging with patient-identifiable data, such as that generated by clinical imaging. The diversity of file formats supported prevents easy visualisation of data without format conversion, and so image data content cannot be viewed in-browser, except for some 2D images and selected accessions where thumbnails have been generated. The archive also does not yet provide a facility to query for specific subcollections (e.g. substantially complete “reference” datasets such as <u>S-BIAD</u>3, *Identification of long noncoding RNAs in regulation of cell division^20^* or <u>S-BIAD4</u>, *A 3D molecular atlas of the chick embryonic heart*^21^). Addressing these limitations are part of the archive’s development plans.

### Planned developments

One of the BioImage Archive’s core aims is to maximise the reuse of imaging data. To use the data, it must first be located. Rich metadata accompanying the data will be increasingly necessary to support data search and comparison^22^. The recently published REMBI recommendations^23^ provide draft metadata guidelines to cover different modalities in biological imaging. The BioImage Archive will implement REMBI, while ensuring that submitters can still deposit data rapidly. Several options exist to support this process, including automated metadata extraction (“harvesting”) and integration with local data-management solutions such as OMERO^24^. We will also integrate emerging community standards, such as the recently proposed 4DN-BINA extensions to the existing OME metadata model^25^. Where possible we will support importing metadata from external applications, such as tools designed to capture image-acquisition metadata at the time of imaging^26^.

EMPIAR provides a useful model for the development of a successful imaging resource. EMPIAR’s early successes came from delivering a valuable service (archiving of raw cryo-EM data) for a focussed and supportive community. EMPIAR’s link with EMDB^27^ allowed depositions to be easier, since most metadata are collected by EMDB, and provided users already familiar with deposition to EMDB with a similar interface and experience. From this beginning, initial use by the core community spread throughout those using cryo-EM and cryo-ET, eventually leading to rapid growth in depositions and requests to support a wider range of data types. To follow this path, the BioImage Archive is working with community networks such as Euro-BioImaging (https://www.eurobioimaging.eu), BioImaging North America (https://www.bioimagingna.org) and Global BioImaging (https://globalbioimaging.org), as well as publishers, to raise awareness, and support data management planning for imaging projects with the aim of guiding data towards the archive on publication.

Supporting the interactive exploration, visualisation and reuse of large imaging datasets is a key long-term goal for the BioImage Archive. This aim is difficult to meet when data are represented across many different file formats and file sizes, and this heterogeneity also presents a significant barrier to reuse. To overcome this barrier requires providing access to data in file formats that are standardised, scalable and optimised for large-scale distribution^28^, in particular by supporting API-based random access to, and downloads of, subsets of data as well as parallel access. At least initially, forcing data depositors to engage in complex data conversion will discourage submissions; therefore, format conversion will need to be carried out within the archive. If community adoption of new file formats grows, the archive can work with commercial imaging system vendors to add support for these formats in their acquisition control and analysis software systems.

Modern deep-learning-based AI techniques have brought rapid progress to the analysis of biological images^29,30^. These technologies often rely on large corpora of reference data, including ground-truths and other annotations to train or retrain models. The BioImage Archive has great potential to accelerate developments in this field by providing such reference datasets and annotations to the community. This requires work to define suitable common formats for annotations, develop deposition pipelines, and encourage and support the community in their use.

Finally, the BioImage Archive will work to grow the ecosystem of added-value databases, by identifying technical or biological domains where specific communities can have their needs served, as well as supporting easy integration for existing or emerging resources. Where large scale image data generation activities, such as cell atlases, need to establish continuous data-deposition pipelines, these can be planned and developed as part of the overarching project. In the long term, the BioImage Archive will need to work globally with the international bioimaging community to establish common standards and distribute data across multiple locations^31^. Similar models involving international consortia support large-scale archives like the ENA and PDB^32^.

### Sustainability and governance

The infrastructure underlying the BioImage Archive is funded by the UK Research and Innovation Strategic Priorities Fund, while staff are funded by EMBL. EMBL-EBI is committed to maintaining the BioImage Archive as an essential part of the collection of EMBL-EBI resources. This is critical for the continued growth and operation of added-value resources (in particular, IDR and EMPIAR) that use the archive as a foundation to support their separate funding efforts. In this way, the archive functions as the basis for a growing bioimaging data ecosystem^14^. As of 2022, archive development is governed jointly through UK Research and Innovation and EMBL-EBI (https://www.ebi.ac.uk/bioimage-archive/project-developments), and receives community input and feedback through its Imaging Ecosystem working group (https://www.ebi.ac.uk/bioimage-archive/project-developments/ecosystem/).

### Conclusion

The challenges of scaling a biological image archive with a very wide scope are considerable. The wide range of different technologies and approaches that together comprise biological imaging gives rise to significant heterogeneity in standards and practices across different communities. This gives rise to a wide diversity of data types, file formats and metadata requirements. Through careful selection of the right level of metadata, integration with local data-management solutions and adoption of new developments in image file formats, we hope to meet these challenges.

A broad repository of open-access FAIR^3^ image data has huge potential to accelerate research in the life sciences. We envision a future in which the BioImage Archive anchors a wide range of community resources representing multiple biological and technical domains, supporting curation and reuse. Together, these would unlock the potential in existing image data, making the investment in biological imaging truly bear fruit.

## Materials and Methods

### Data access

Access to images and other files is provided both directly through the web portal, and via an API (https://www.ebi.ac.uk/biostudies/help#API). Images can be downloaded directly from the web portal via the HTTPS protocol, as well as via FTP or Aspera, while Globus support is planned. The API supports programmatic access to both data and metadata. This allows enumeration of datasets, keyword and advanced search, retrieval of individual image-level metadata and download of individual files.

### System architecture

The three main components of the system are its backend that provides data and metadata management and lifecycle services, the submission tool that enables dataset deposition, and the data-access interface for both web and programmatic access.

The BioStudies platform implements a very generic information model, permitting hierarchical metadata annotations, attaching and annotating groups of data files to various points in the dataset hierarchy, and using the concept of external links to point to related components of data in other resources. A submission to the BioImage Archive invokes an imaging data-specific web form, guaranteeing that the data submitted in this manner will have useful metadata attributes defined. The submission tool uses the Angular web application framework, and is written in TypeScript.

When submission is completed, files are uploaded to FIRE (FIle REplication), EMBL-EBI’s very-large-scale object data storage system. This provides geo-dispersed long-term sustainable storage, operational redundancy, and backup to tape. Dataset level metadata are stored in a MongoDB database. The system backend is coded in Kotlin.

The metadata database is indexed nightly to power the search engine that is a part of the data access application and exported to allow distributed data hosting. The search engine is built using Apache Lucene, and the data access system is a Java web application.

## Acknowledgements

We thank all of the community representatives that participated in the workshops that established the scientific rationale for the BioImage Archive and the REMBI metadata framework. We also thank the IDR, EMPIAR, BioStudies and BioImage Archive teams, EMBL-EBI’s Bioimaging project team, Operational and Capital Projects team, and the storage team within EMBL-EBI’s Technical Services Cluster. Finally, we would like to thank all the scientists, past, present and future, who deposit their data in the BioImage Archive.

## Funding statement

The development of the BioImage Archive has been supported by European Molecular Biology Laboratory member states, Wellcome Trust grant 212962/Z/18/Z and UKRI-BBSRC grant BB/R015384/1. Work on EMPIAR was funded from 2014 to 2021 by two project grants awarded to EMBL-EBI by UKRI-MRC and UKRI-BBSRC (MR/L007835/1 and MR/P019544/1). From 2021, EMPIAR benefits from funding from the Wellcome Trust (221371/Z/20/Z). The EMBL-EBI IT infrastructure supporting the BioImage Archive is funded by the UK Research and Innovation Strategic Priorities Fund.

## Notes

### Competing Interest Statement

The authors have declared no competing interest.

### Summary of Updates

This version of the manuscript has been revised to improve overall readability and clarity. Figure 1 has been made clearer, Figure 2 now includes more data and an explanatory table has been added.

